# Metachronal wave coordination encodes multimodal swimming in ciliated unicellular predators

**DOI:** 10.1101/2025.09.12.675801

**Authors:** Aikaterini M. Kourkoulou, Maggie Liu, Arnold J. T. M. Mathijssen, Guillermina R. Ramirez-San Juan

**Affiliations:** Institute of Physics, EPFL, Lausanne, CH 1015; Department of Physics, University of Pennsylvania, Philadelphia, PA 19104

## Abstract

Motile cilia are slender cellular appendages, conserved across eukaryotes ranging from unicellular protists to humans, that beat to generate fluid flow. In most organisms, cilia form dense arrays of thousands of filaments that coordinate their motion into persistent, temporally synchronized patterns known as metachronal waves. Despite their ubiquity, the dynamics of these patterns and their role in tuning propulsion remain poorly understood. Here, we investigate how metachronal coordination shapes the navigation of *Didinium nasutum*, a highly agile unicellular predator with two circumferential ciliary bands. Using high-speed imaging of freely swimming cells, we capture and quantify the dynamics of metachronal wave coordination and track their evolution across different swimming states and transitions. Combining these measurements with a hydrodynamic model, we uncover how dynamic changes in coordination directly regulate propulsion and maneuverability. We show that stable metachronal waves support persistent directed swimming, local inhomogeneities in coordination give rise to curved trajectories, and global wave reversals accommodate rapid evasion-like reorientations. Our findings reveal that transitions between coordination modes allow *Didinium* to access a diverse swimming repertoire, highlighting dynamic ciliary patterning as a key mechanism to encode complex microscale navigation strategies. More broadly, they provide mechanistic insights into how metachronal coordination shapes fluid flows generated by dense ciliary arrays found in unicellular protists and airway epithelia alike, ultimately influencing swimming and transport in biological systems.

---

The ability to generate flows with spatial and temporal precision is critical for the function of diverse biological systems, including mucus transport in mammalian airways, cerebrospinal fluid circulation within brain ventricles, and propulsion in motile unicellular organisms [1–3]. In each of these contexts, flow is driven by slender hair-like organelles that extend from the surface of eukaryotic cells, known as cilia. These organelles beat in periodic asymmetric wave patterns, allowing effective fluid movement at the microscale. In most biological contexts, thousands of cilia are organized in dense arrays and coordinate their motion in time, forming metachronal waves (MWs), a striking example of collective behavior in which each cilium beats with a constant phase difference relative to its neighbors [4–6].

Theoretical and computational studies have shown that metachronal coordination provides key functional advantages to ciliary arrays, including improved fluid transport and pumping efficiency [7–9]. Recent experimental work has begun to uncover the dynamics of metachronal waves in living organisms, characterizing their emergence, directionality, and stability in ciliated tissues and microswimmers [10–13]. However, the relationship between the spatio-temporal dynamics of MWs and their biological function remains poorly understood. Studies of unicellular algae bearing a small number of cilia have revealed that shifts in their temporal coordination underlie abrupt shifts in cell motility, enabling rapid behavioral responses. For instance, in the green alga *Chlamydomonas reinhardtii*, spontaneous transitions between synchronous and asynchronous beating of its two cilia allow the cell to rapidly reorient in response to environmental stimuli [14–16]. Similarly, in multiflagellate algal species, dynamic modulation of phase differences between neighboring cilia enables access to distinct motility gaits, allowing the cell to control swimming direction and speed in absence of a nervous system [17, 18]. In the case of dense ciliated carpets on the surface of many protists, changes in metachronal coordination, such as cilia and MW reversals, have been presumed to underlie cell reorientation and backward swimming [19–21]. However, the kinematics of MWs during these transitions have never been directly measured, limiting our understanding of how metachronal wave dynamics regulate adaptive cell navigation.

To gain insight into the functional relevance of metachronal waves, here we study how temporal control of metachronal coordination gives rise to a wide repertoire of swimming behaviors in the ciliate *Didinium nasutum*. This unicellular organism has a nearly spherical body and thousands of cilia arranged in two circumferential bands oriented perpendicular to its anterior–posterior axis [22, 23] (Fig. 1A). This distinct morphology enables direct visualization of metachronal coordination throughout the entire ciliary array (Fig. 1B,C). *Didinium* is a predatory ciliate that uses a specialized proboscis to capture fast swimming prey, mainly *Paramecium* [24, 25]. As a highly agile predator, it swiftly shifts between a variety of swimming patterns, including straight or helical motion, rapid reorientation, and spinning on surfaces (Fig. 1D, Movie 1). Its ciliary arrangement and rich swimming dynamics position it as an ideal model for investigating how metachronal coordination encodes swimming robustness and flexibility, enabling complex motility.

**FIG. 1.**
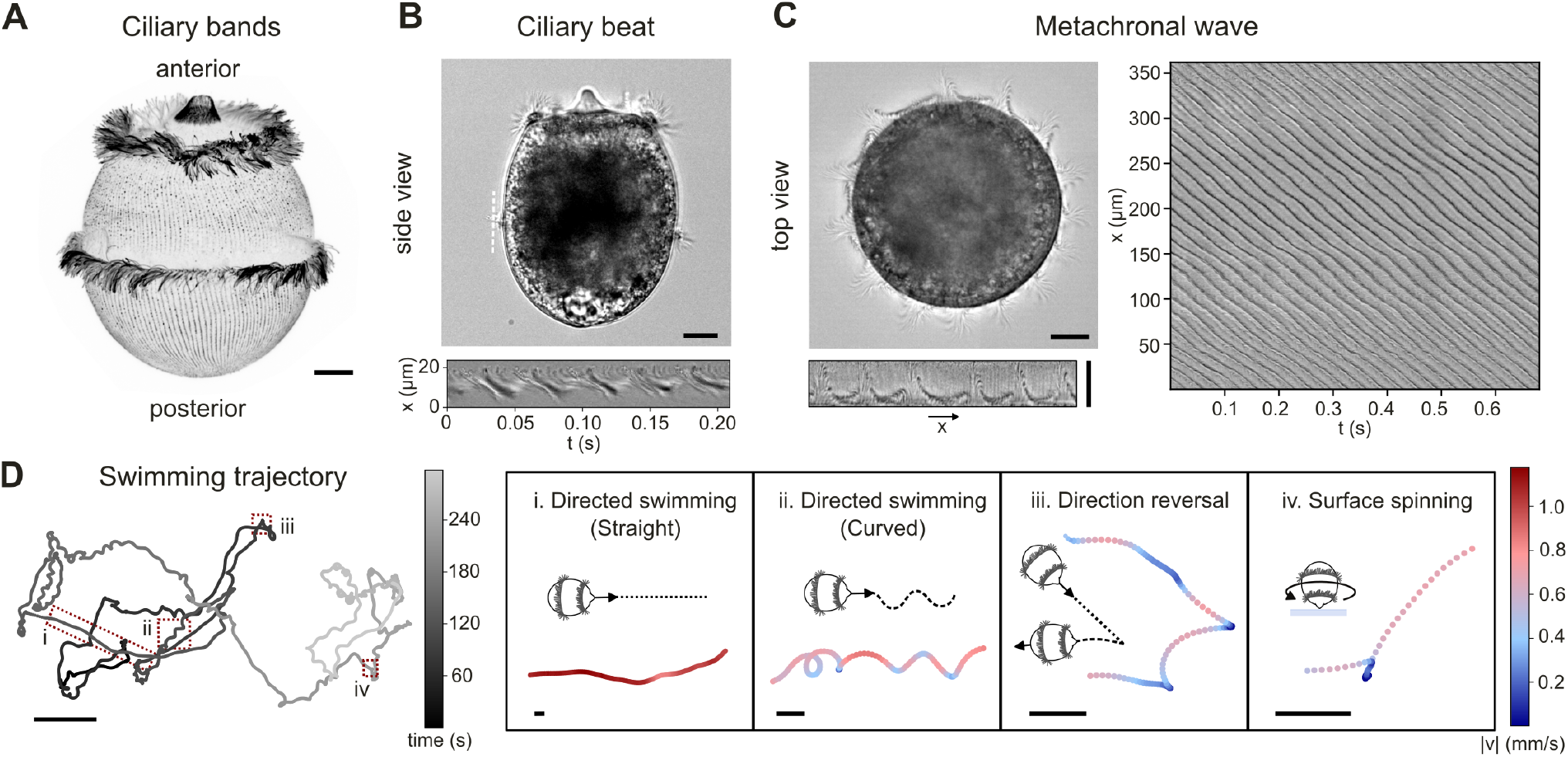
The spatiotemporal cilia patterning of *Didinium nasutum* supports diverse swimming behaviors. (A) Immunoflorescence image (poly-E) of the localization of cilia in *Didinium nasutum*. The cilia are organized in two dense ciliary bands at the anterior and the midsection of the cell. (B) DIC microscopy image of a transverse view of a free-swimming cell. The dashed white line marks the axis along which the kymograph (bottom) was extracted, visualizing the ciliary beat pattern over time. (C) Left, top: DIC microscopy image of an orthogonal (top-down) view of the cell. A metachronal wave propagates across the full diameter of the ciliary array. Left, bottom: A segment of the ciliary band after polar transform, showing individual wavecrests of the MW. Right: Kymograph of intensity variations along the cell circumference, showing persistent propagation of the metachronal wave along the cell circumference. (D) Swimming trajectory of a single cell. Red dashed rectangles mark trajectory segments highlighted in the insets on the right, each illustrating a distinct type of motion over a 4 second interval: (i) directed fast swimming along a straight path, (ii) directed fast swimming along a curved path, (iii) rapid reversals in swimming direction, and (iv) spinning on a surface without net translation. Scale bars: 20 µm (A-C), 2 mm (D) and 200 µm (insets in D).

In this work, we employ high-speed imaging of free-swimming cells to directly link dynamic ciliary coordination with cell motility, revealing how distinct ciliary patterns control specific motility states and transitions. We find that stable dexioplectic metachronal waves give rise to persistent straight swimming, while local spatial pattern inhomogeneities result in helical and curved trajectories. We further demonstrate that cells are capable of rapid reorientations, controlled by global wave reversals that propagate along the ciliary bands, enabling precise changes in direction reminiscent of escape responses. To gain a deeper understanding of these mechanisms, we develop a computational model that replicates cilia-driven force generation under different coordination regimes, allowing us to test how these patterns shape cell motility and steering. Finally, we demonstrate that rapid transitions between different coordination modes allow cells to access a broad and diverse swimming repertoire, highlighting the central role of dynamic ciliary coordination in encoding complex microswimmer behavior.

## GLOBAL DEXIOPLECTIC COORDINATION DRIVES STEADY STRAIGHT MOTION

We first characterized the cell geometry and ciliary organization using immunofluorescence microscopy (see Methods § 1 b). Consistent with previous observations, we find that cells possess two circumferential ciliary bands oriented perpendicular to the anterior-posterior (A-P) axis: one located anteriorly, just below the proboscis, and the other positioned in the midsection of the cell (Fig. 1A). Although the length of the A-P axis can very slightly between cells (122 ± 9 µm), the cell body maintains a consistent aspect ratio (1.12±0.07). Furthermore, the position of the two bands remains conserved between individuals, as indicated by a constant ratio between band separation and the length of the A-P axis (Fig. S1).

Given that cells can abruptly change their swimming pattern, we hypothesized that the observed diversity of gaits arises primarily from variations in the beating dynamics and coordination of the ciliary bands. To explore this, we investigated the temporal coordination of ciliary beating within the two bands using ultra-fast imaging of freely swimming cells (see Methods § 1 c). During directed straight swimming, cells move forward while spinning about their A-P axis. Cilia beat parallel to the cell axis from anterior to posterior with an off planar waveform (Fig. 1B, Movie 2) and metachronal waves propagate along the circumference in both ciliary bands, traveling in the same direction as the cell spin (Fig. 2A, Fig.S2A, Movie 3). During this motility state, cells are oriented parallel to the field of view, limiting observation of metachronal waves to a subset of the ciliary bands. Interestingly, *Didinium* has a natural tendency to stably spin against interfaces, and is often found spinning in place with its anterior end against the bottom of the coverslip. In this orientation, each ciliary band lies approximately in a single plane, allowing global wave coordination to be tracked over time by means of intensity kymographs (Fig. 1C, Movie 4). Notably, the direction of the propagating wave for both motility states is always dexioplectic (counterclockwise when viewed from the anterior end, with cilia power stroke from anterior to posterior) in agreement with classical studies [26]. We note that in the case of stationary spinning, coordination persists in the medial band; however, the anterior band appears uncoordinated even though the cilia are beating (Fig. S2B, Movie 5). Further analysis of this state is thus restricted to the medial band.

**FIG. 2.**
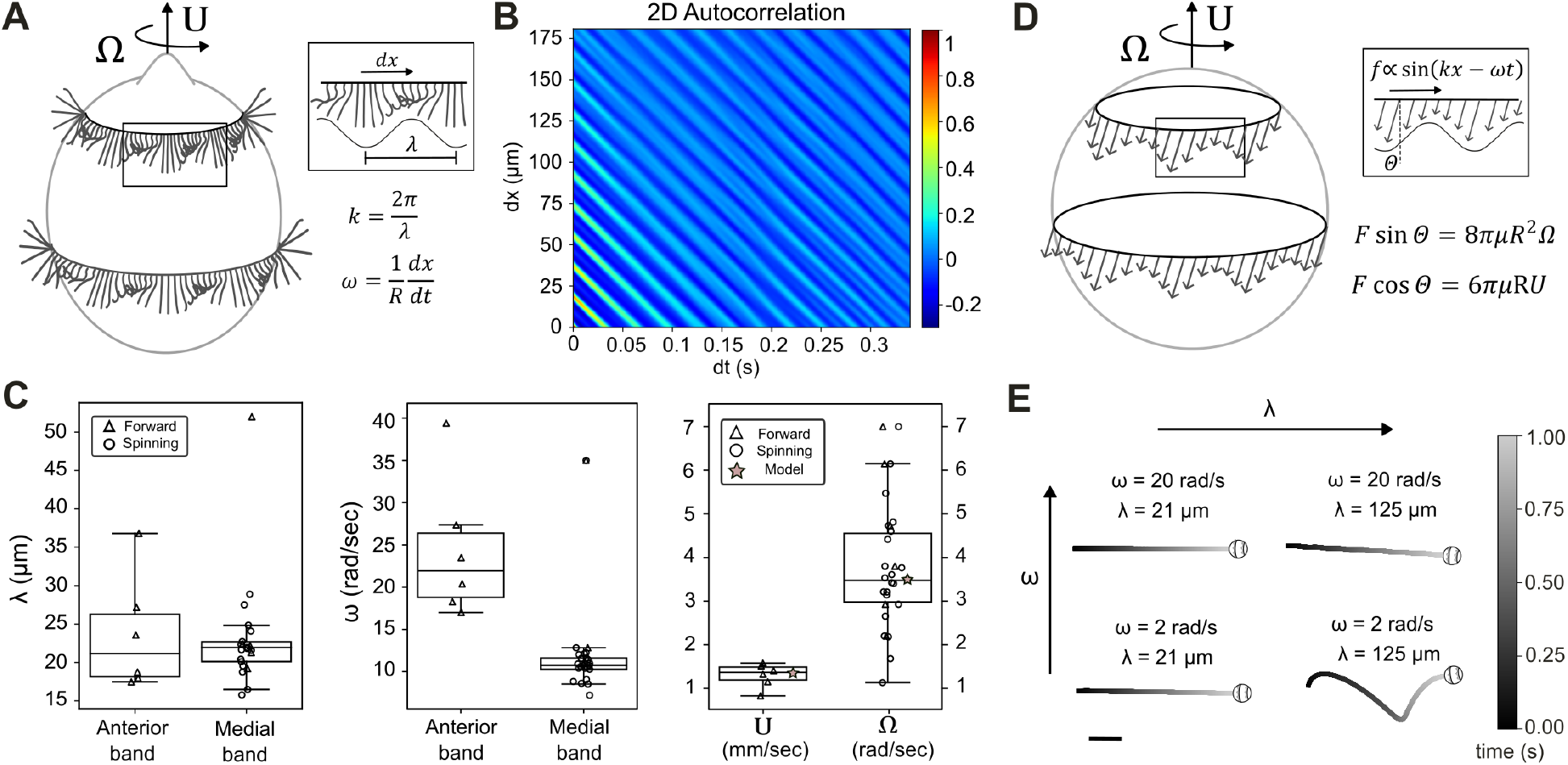
Persistent dexioplectic metachronal waves around the cell circumference support straight swimming. (A) Schematic of the dexioplectic MW on the two ciliary bands. Cell translational velocity, *U*, and spinning velocity, Ω, are indicated by arrows. Inset: Enlarged view highlighting the main MW parameters. (B) Two-dimensional autocorrelation of intensity along the medial band, showing global persistent coordination. (C) Metachronal wavelength (left) and wave velocity (middle) of the anterior and medial ciliary bands during forward swimming and surface spinning. Cell translational and spinning velocity (right) for the same conditions. Mean values were used to parametrize the model. N_*f*_ =6 forward-swimming cells, N_*s*_ = 16 surface-spinning cells. (D) Schematic of the computational model. Ciliary bands are represented as an array of point forces with a sinusoidal envelope propagating along the circumference. Inset: Enlarged view and definition of the forces amplitude and direction. (E) Simulated trajectories for homogeneous metachronal waves with varying wave velocity and wavelength. Trajectories remain straight except under extreme conditions of long wavelength and low angular velocity, where the pattern loses symmetry. Scale bar: 200 µm.

Studying free-swimming cells allowed us to measure cell motility and metachronal wave properties simultaneously (Extended data Table I). Within the observed time windows, forward swimming cells move with stable high linear velocities equal to approximately 11 body lengths per second (*U* = 1.3 ± 0.25 mm/s) following approximately straight trajectories. Similarly, cell spin remains stable during both directed swimming (Ω = 4.5 ± 1.7 rad/s) and surface spinning (Ω = 3.3 ± 1.1 rad/s), as long as metachronal wave propagation is maintained (Fig. S2D). To measure the MW parameters, we calculate the two-dimensional autocorrelation of the image intensity along each ciliary band for both of these motility states (see Methods § 2 b). Metachronal patterns remain robust both in time and space and demonstrate global phase locking (Fig. 2B). We find that the wavelength is comparable for both bands across both types of motion (*λ* = 24 ± 7 µm for directed swimming and *λ* = 22 ± 3 µm for surface spining)(Fig. 2C). However, during directed forward swimming the anterior band cilia beat at a higher frequency (35.5 ± 2.2 Hz), compared to the medial band (28 ± 3 Hz) (Fig. S2C), resulting in a higher wave propagation velocity and suggesting that the anterior band plays a more dominant role in force generation and propulsion.

To gain a mechanistic understanding of how metachronal wave patterns affect propulsion and shape cell trajectories, we built a computational model that simulates the force generation profile of the two ciliary arrays. Each band was represented as a sinusoidal array of forces following the experimentally measured wave direction and velocity, with individual force vectors oriented at an angle, Θ, relative to the swimming direction to capture the tangential component responsible for cell rotation (Fig. 2D). In this reduced model, the swimming and rotational velocities of the cell, *U* and Ω, are determined by the balance of hydrodynamic forces, governed by the Stokes equation (see Methods § 3). By fitting the model to the experimental swimming and rotation speeds, we estimate a net propulsive force of *F* = 1.3 · 10^−9^*N* and Θ = 14°, consistent with previous literature [26, 27].

By varying the MW parameters in the simulation, we find that cell trajectories remain straight across a wide range of wavelengths and propagation speeds (Fig. 2E).

This suggests that metachronal waves play a key role in stabilizing cell orientation and ensuring robust, directed motion, with short wavelengths producing a more uniform force distribution that suppresses small trajectory oscillations and higher propagation speeds further enhancing swimming stability. Pronounced deviations from straight paths arise only at low propagation speeds and very large wavelengths where the sparse wavefront distribution breaks the inherent symmetry of the force field (e.g. N=3 periods along the circumference). Since such conditions lie outside the experimentally observed MW parameter range, curved trajectories must originate from additional asymmetries in force generation rather than from simple modulation of metachronal wave parameters.

## METACHRONAL WAVE INHOMOGENEITIES GIVE RISE TO CURVED SWIMMING PATHS

During steady straight swimming, the average net force generated by the ciliary arrays remains homogeneous and constant. However, in addition to linear trajectories, cells frequently exhibit helical swimming paths with varying geometric parameters (Fig. 3 B,C). For a cell to swim in a curved trajectory, there must be an asymmetry in the spatial distribution of propulsive forces, that generates a net torque changing the swimming direction (Fig. 3A) [16, 28]. To investigate potential sources of asymmetry in propulsive forces, we analyzed metachronal wave dynamics by observing the medial band during inplace spinning, which offered a global view over extended timescales that is not possible to obtain during free directed swimming.

**FIG. 3.**
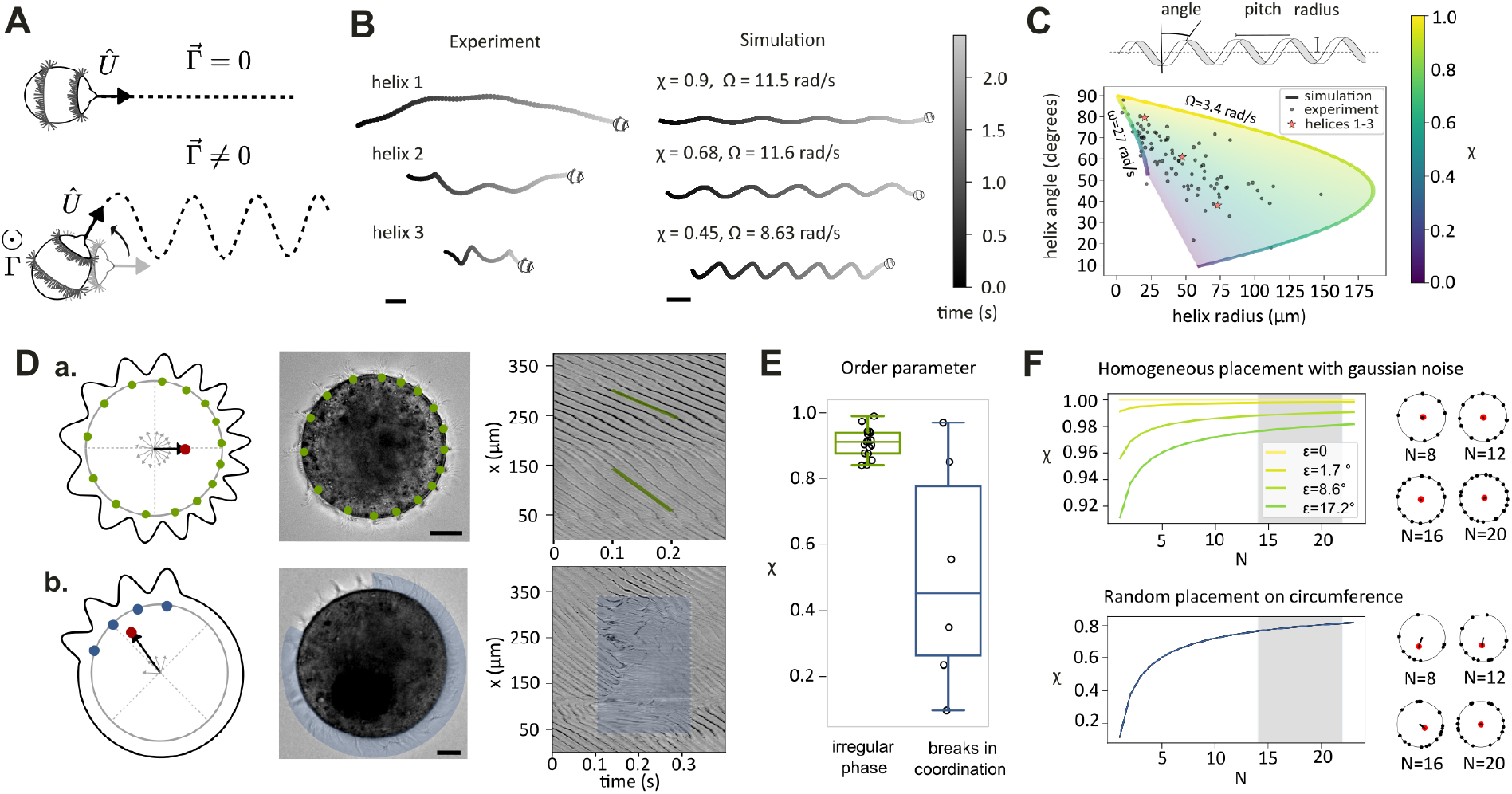
Torques generated by MW inhomogeneities control curved paths. (A) Homogeneous propulsion fields produce straight trajectories (top), whereas force heterogeneity generates torques that lead to helical paths (bottom). (B) Left: Examples of experimental helical trajectories of the same cell at different time points. Scale bar: 200 µm. Right: Simulated trajectories fitted to the experimental examples, showing that different degrees of heterogeneity, quantified by 𝒳 are required. Scale bar: 500 µm. (C) Top: Schematic defining the main geometrical parameters of a helix. Bottom: Helix angle as a function of helix radius for experimental helical segments (N = 93 from five cells) compared with model predictions varying Ω and 𝒳 (shaded area). Bold lines indicate the upper and lower limits. Red stars mark the examples from panel B. (D) Asymmetry in force distribution. (a) Left: Schematic and DIC image illustrating wavecrest positions (green dots) for a cell with nonuniform phase distribution. Black arrow indicates the resulting center of force (red dot). Right: Kymograph of intensity along the circumference, with green lines showing different wave velocities in two regions. (b) Left: Schematic and DIC image of a cell with partial loss of coordination, showing wavecrest positions (blue dots). Right: Kymograph marking the uncoordinated region (blue shaded area). Scale bars: 20 µm. (E) Boxplots of the degree of heterogeneity, 𝒳, in the medial band of spinning cells, comparing phase irregularities (green, N_*a*_ = 16) and loss of coordination (blue, N_*b*_ = 6). (F) Scaling of 𝒳 with the number of wavecrests *N*. Curves represent a CLT-based approximation. Points are either perturbed from uniform placement with Gaussian noise (top) or randomly distributed (bottom). Shaded areas indicate the experimentally measured range of *N*. Right: Representative schematics showing wavecrest positions (black dots) and the resulting center of force (red dot) for increasing *N*.

Our analysis revealed two distinct mechanisms capable of generating such steering torques. First, we observed that the phase difference between neighboring cilia can vary along the circumference of the ciliary bands. This variation creates an asymmetry in the wave pattern, producing broader wavelengths and faster-traveling wavefronts on one side of the cell (Fig. 3D.a, Movie 6). A second source of spatial asymmetry comes from temporal disturbances in metachronal coordination, where local breaks are observed in a subset of the ciliary band, while MWs continue to propagate in the rest of the circumference (Fig. 3D.b, Movie 7). The life time of these “glitches” in coordination is short (0.14 ± 0.08 s), and the size of the affected region can vary significantly from 25 to 300 µm corresponding to 6-72% of the cell circumference (Fig. S4C). We observe that both of these inhomogeneities are fixed in the reference frame of the cell and therefore travel around the A-P axis at the same velocity as the cell spin (Movies 6,7).

To quantify the degree of heterogeneity, we treated each wavefront of the metachronal wave as an individual oscillator and calculated the complement of the Kuramoto order parameter: 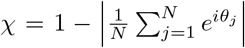, which ranges from 0 (disorder, maximum asymmetry in wave-front placement) to 1 (order, homogeneous distribution of wavefronts) [29]. Only the wavefronts at the regions where the metachronal wave remained coherent were considered.

We quantified 𝒳 for 16 individual spinning cells with wavelength asymmetries and measured a range of 0.84 to 0.99 (Fig. 3E, irregular phase). Typically, each cell maintained a stable 𝒳 throughout the observation window. However, we occasionally observed instances where an individual cell, after brief local loss of coordination, switched to a different 𝒳 value which was subsequently stably maintained, indicating that cells can transition between different degrees of asymmetry. In the case of coordination glitches, the larger variability in size resulted in a broader 𝒳 distribution ranging from 0.1 to 0.9 (Fig. 3E, breaks in coordination).

To test whether these mechanisms could generate the torques needed to explain the diversity of swimming trajectories observed, we used our model to predict the helix characteristics across the observed range of 𝒳 by modifying the force envelope, allowing for local loss of coordination (where forces are zero), as well as an asymmetric profile of phase shift, akin to the experimentally observed phenotypes (Fig. 3D, Fig. S5). While the baseline is set so that these inhomogeneities travel with the spinning velocity of the cell, Ω ∼ 3.5rad/s (upper border), we additionally explored a broader range, *ω* = 3.5 to 27 rad/s (Fig. 3C shaded region), corresponding to the MW angular velocities. We conclude that for this range, and 𝒳 in the range 0-1, over 93.4% of experimentally measured data fall within the analytical prediction. Thus, the observed variations in MW wavefront coordination are capable of generating sufficiently strong torques to account for the observed trajectories.

To systematically investigate how variations in wavefront number and placement influence asymmetry and trajectory stability, we applied the central limit theorem to derive analytical expressions for 𝒳 as a function of the number of wavefronts *N* under two scenarios: (i) a nearly uniform distribution of wavefront with tunable Gaussian noise, and (ii) a completely random arrangement of wavefronts (Fig. 3F). Our analysis shows that while noise in an otherwise uniform distribution only weakly perturbs trajectory helicity, random wavefront placement through irregulated wavelengths can induce substantial asymmetry. However, for both cases increasing the number of wavefronts systematically suppresses asymmetry, stabilizing swimming trajectories and promoting straighter paths. We additionally note that the experimentally observed range of wavefront numbers falls near the plateau of the curves, where 𝒳 is relatively insensitive to *N* variations, indicating an inherent robustness in the system’s swimming dynamics.

Altogether, these results show that by harnessing both local differences in metachronal wave dynamics and the collective organization of wavefronts, the cell gains access to a wide variety of swimming trajectories while maintaining robust control, highlighting how dense ciliary bands enable precise, efficient control of global motil-ity through minimal local changes.

## CILIA AND METACHRONAL WAVE REVERSALS LEAD TO TUNABLE DIRECTIONAL CHANGES

One well-known behavioral response in many ciliates is the avoidance reaction; a rapid, stimulus-triggered reversal in swimming direction that allows cells to escape unfavorable conditions. Typically induced by mechanical or chemical cues, this response has been primarily studied in *Paramecium*, where a reversal in the direction of the ciliary effective stroke drives backward motion [21, 30]. In most cases, such changes in ciliary dynamics are inferred indirectly through flow tracking or cell trajectories rather than directly resolving the underlying ciliary or metachronal wave reconfigurations [19]. In the case of *Didinium*, similar rapid directional reversals occur spontaneously in free-swimming cells (Fig. 4A), allowing us to measure directly the dynamics of ciliary motion and metachronal wave changes during these events.

**FIG. 4.**
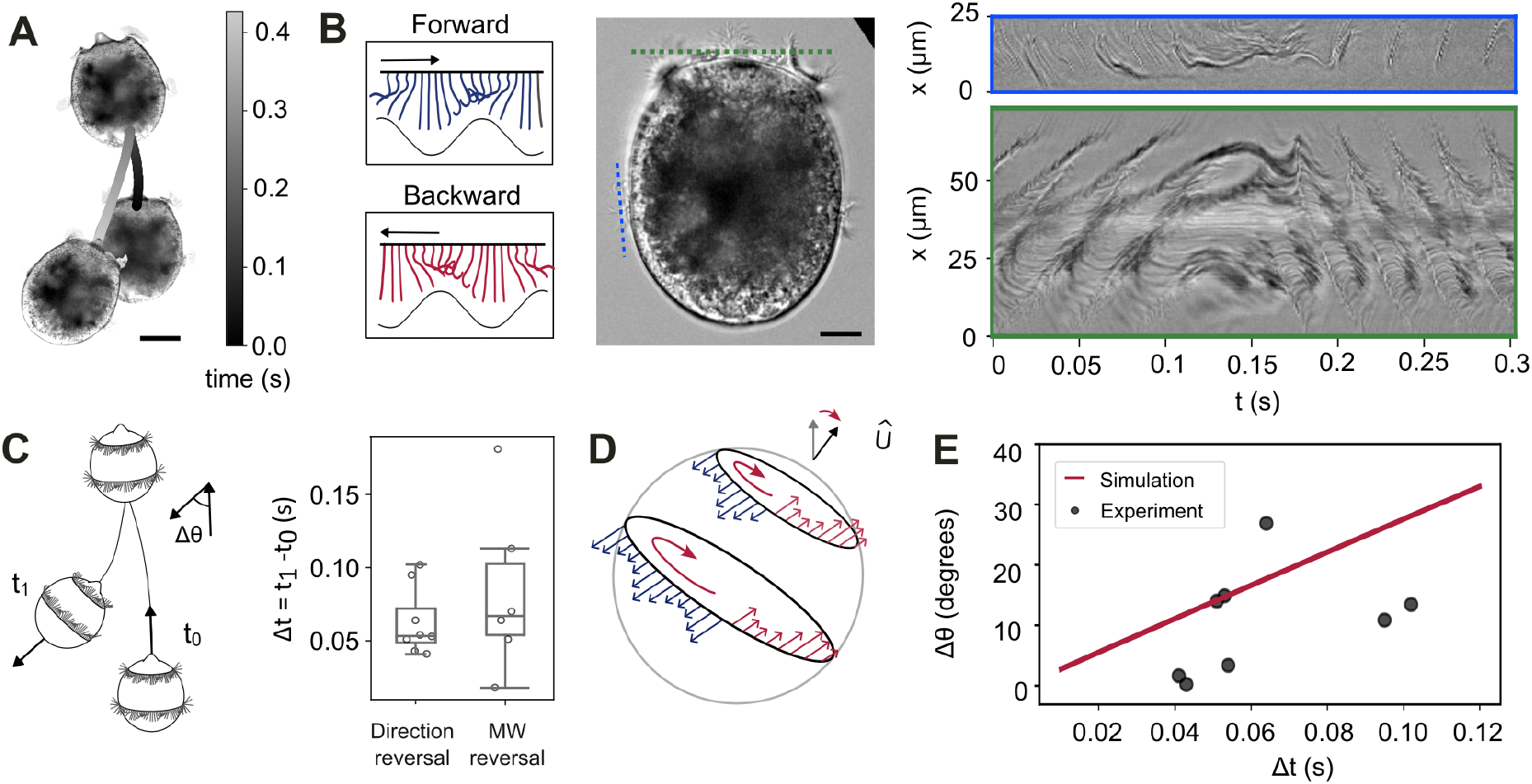
MW reversals drive sharp cell reorientations. (A) Trajectory of a cell undergoing a sharp reorientation from forward to backward swimming. DIC microscopy snapshots show the initial orientation, a transient immobilization, and the final orientation. Scale bar: 50 µm. (B) Left: Schematic illustrating reversal of metachronal wave coordination before (blue) and after (red) the transition. Right: Snapshot from a registered timelapse of a reorienting cell. Blue and green lines indicate the regions used to generate the kymographs (right). During reorientation, the effective ciliary stroke reverses (blue window), and the MW also switches direction after a brief pause (green window). Wave reconfiguration occurs progressively rather than simultaneously as seen by the delay in the reestablishment of the two wavefronts. Scale bar: 20 µm. (C) Left: Schematic representation of the cell axis reorientation. *t*_0_ corresponds to the start of reorientation and *t*_1_ to its end. The total deflection angle is denoted as Δ*θ*. Right: Boxplots of the reorientation duration and MW reversal duration. (D) Simulation of MW reversal, implemented by sequential inversion of force direction. During the transition, original (blue) and reversed (red) waves coexist in different regions of the array until the reversal is complete. (E) Deflection of the cell axis after rapid reorientation as a function of time. Line indicates the model prediction and scatter points correspond to experimental measurements. *N*_*cells*_=8.

By tracking individual cells (see Methods § 2 c), we find that reversals from forward to backward swimming occur very rapidly, typically completed within 0.06 ± 0.02 s. During these events, the swimming direction changes only slightly, with deflection angles ranging from 0° to 26° (Fig. 4C). By analyzing high-speed time-lapse recordings, we resolve both ciliary activity and metachronal coordination during these transitions. We observe that, coincident with the reorientation of the swimming direction, the cilia temporarily halt their beating before resuming with a reversed effective stroke (Fig. 4B, top blue boxed kymograph, Movie 8). Simultaneously, individual metachronal wavefronts appear to pause, then resume traveling in the opposite direction (Fig. 4B, bottom green boxed kymograph, Movie 8). The timescale of the reorganization in ciliary coordination (0.08 ± 0.05 s) closely matches the time it takes for the cell to complete its directional reversal (Fig. 4C). Notably, the reversal in wave propagation does not happen all at once across the entire cell surface but appears to occur gradually along the ciliary band (Fig. S6A).

To assess the impact of the observed reconfigurations on cell motility, we use our model to simulate MW beating pattern reversals and compare the resulting paths with those observed experimentally. We model the metachronal wave reconfiguration as a locally triggered reversal of ciliary beating that propagates linearly around the circumference: once a group of cilia reverse their beat direction, neighboring cilia follow in succession, gradually inverting the original pattern. The direction of the sinusoidal envelope corresponding to the inverted region is simultaneously reversed (Fig. 4D, Movie 9). We find that during this process, the cell gradually reorients, resulting in a final deflection angle that increases linearly with the duration of the MW beating pattern rearrangement, marginally exceeding the values observed experimentally (Fig. 4E). Collectively, our results show that the gradual and propagating nature of wave reversals, rather than a simultaneous switch across the whole ciliary band, enables the cell to access a tunable range of swimming deflections, with potential significance in evasion and hunting strategies.

## TRANSITIONS BETWEEN WAVE PATTERNS ENABLE A BROAD TRAJECTORY SPACE

After identifying the distinct metachronal patterns that give rise to specific motility gaits, we sought to determine how the transition between these states shape swimming behavior over extended timescales. To this end, we analyzed long-term swimming trajectories to capture the full spectrum of behavioral variability. We observed velocity and orientation patterns that qualitatively match the previously described motility gaits, and used these patterns to classify trajectories into distinct states (Fig. 5A, see Methods § 2 d).While each state exhibits stereotypical features, the cell’s velocity and flexibility, quantified by the angular velocity, span a broad range of values (Fig. 5B). By analyzing the duration of each state through survival probabilities, we find that while the spinning state can typically persist for many minutes, directed swimming is often transient, and frequently interrupted by rapid reorientation events (Fig. 5C). Together, these rapid state transitions and the broad range of explored velocities and orientations enable the cell to continuously navigate a highly flexible and dynamic swimming space, highlighting the sophisticated control of motility that arises from local coordination of its ciliary bands.

**FIG. 5.**
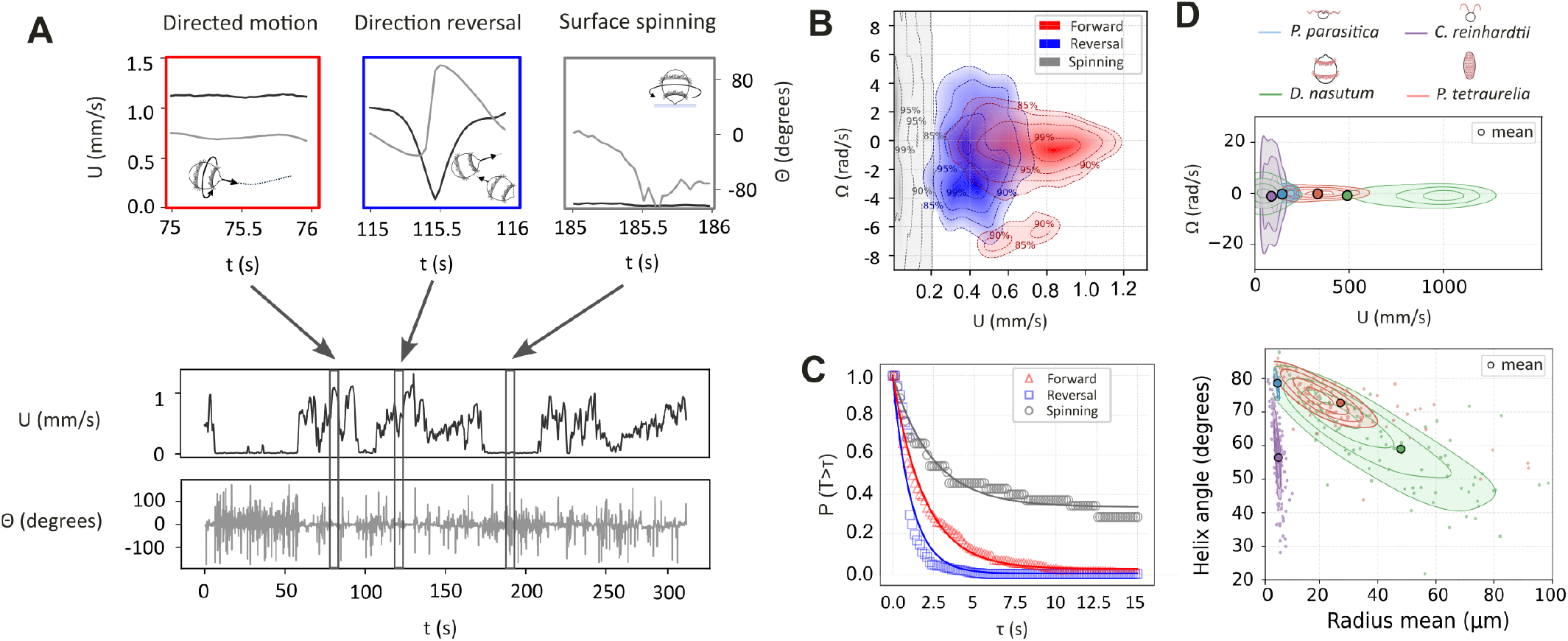
Switching between metachronal wave states enables diverse swimming behavior. (A) Top: Linear velocity (black) and cell orientation (gray) over time, illustrating three distinct motility states: directed swimming (red), rapid directional reversals (blue), and surface spinning (dark gray). Bottom: Linear velocity (black) and orientation (gray) over a 5-minute trajectory. Rectangles indicate the segments corresponding to the insets above. (B) Velocity density maps, generated using kernel density estimates (KDEs). Filled contours represent densities above the 85th percentile. Labeled contours indicate the 85th, 90th, 95th, and 99th percentiles. Each gait is qualitatively stereotypical but spans a broad range of velocity magnitudes. (C) Survival probability of the three motility states. Empty symbols denote experimental values; colored lines show exponential fits (forward swimming: *λ* = 0.511, *R*^2^ = 0.986; reorientation: *λ* = 0.407, *R*^2^ = 0.944; spinning: *λ* = 0.869, *R*^2^ = 0.926). (D) Velocity density maps (top) and helix characteristics (bottom) for four different ciliated swimmers. Shaded contours indicate KDE density levels (40%, 60%, 80%, 95% of maximum). Bold circles mark mean values and transparent points correspond to individual helical segments. Five individuals per species were analyzed (1 minute tracks for the zoospores, 5 minute tracks for other species).

To understand how the specific ciliary arrangement of Didinium shapes its swimming repertoire, we compared its motility with that of other microswimmers. Tracks for *Phytophthora parasitica* zoospores and the alga *Chlamydomonas reinhardtii* were extracted from the published studies [31, 32] and reanalyzed alongside our experimental data for *Didinium* and *Paramecium tetraurelia*. These species were selected to illustrate distinct ciliary strategies: the zoospore and *C. reinhardtii* have relatively few cilia, favoring propulsion suited to long-range swimming or phototactic navigation, whereas *Paramecium* has fullbody ciliary coverage in contrast to the distinct ciliary bands of *Didinium*. We quantified linear and angular velocities as well as helix geometrical parameters to capture the diversity of trajectories each species can adopt (Fig. 5D). *Didinium* spans a wider range of velocities and helix variability than the other species, reflecting how its discrete, dense ciliary bands combine high cilia number for fast propulsion, matching or exceeding the speeds of its prey, with the ability to harness small asymmetries, as in *C. reinhardtii*, to achieve rapid and flexible maneuvering, essential for succesful prey capture.

## DISCUSSION

In this work, we provide a comprehensive analysis of metachronal wave dynamics during free swimming in the predatory ciliate *Didinium nasutum*, demonstrating how temporal and spatial modulation of ciliary coordination underpins the organism’s remarkable swimming versatility. We show that stable, global dexioplectic metachronal waves along the two circumferential ciliary bands generate persistent, high-speed straight swimming. Local inhomogeneities in cilia phase difference and temporary dissolution of coordination give rise to curved and helical trajectories by introducing asymmetries in force distribution, enabling path flexibility. Moreover, we reveal that rapid changes in swimming direction are controlled by a progressive inversion of the cilia beat and metachronal wave direction, allowing for tunable turning angles crucial for effective and rapid maneuvering. Through a computational model parametrized with experimental values, we quantitatively demonstrate how these distinct wave coordination regimes translate into specific swimming gaits, offering mechanistic insight into the functional role of metachronal waves in cell navigation.

In motile ciliated swimmers, cilia are commonly arranged in dense carpets covering the entire cell surface. Alternatively, cilia can be found in single ciliary bands, typically located near the cell’s anterior. In this arrangement, ciliary coordination often facilitates feeding currents, supporting both sessile and motile micro-grazers [33–35]. This spatial organization contrasts with the two distinct circumferential ciliary bands found in *Didinium*, raising the question of whether this unusual architecture provides specific functional advantages for a predator that does not rely on filter feeding.

Our results suggest a functional division of labor between these two bands. The anterior band beats at a higher frequency and generates stronger flows, suggesting it plays a dominant role in forward propulsion. In contrast, the medial band may serve a more modulatory function, potentially contributing to directional stability or enabling rapid reorientation. This spatial separation of ciliary bands may allow the cell to compartmentalize propulsion and maneuvering, with the anterior band generating thrust and the medial band producing torque for fine-tuned control. Additionally, our analysis indicates that small, localized changes within a ciliary band are capable of steering torques sufficient to significantly change cell swimming paths. This is especially advantageous for precise maneuvering during hunting of large, fast-moving prey, and would allow for more rapid reactions than a fully ciliated surface, which would require a larger scale of coordination rearrangement to produce a similar effect.

The ability of *Didinium* to rapidly and locally modulate ciliary coordination suggests that cilia coupling is not merely supported by hydrodynamic interactions. Our observations reveal that cilia on opposite sides of the cell can independently adjust their phase relationships, leading to transient asymmetries and localized disruptions in metachronal wave patterns. When these disruptions repeatedly occur for the same individual cell, they tend to recur approximately at the same region of the cell surface, implying the existence of internal structural asymmetries that bias where coordination can be selectively weakened. The persistence and spatial specificity of these events indicate that fine-scale control over inter-ciliary coupling is likely mediated by intracellular mechanisms such as underlying protein scaffolds, membrane compliance, or localized ion channel distributions that allow spatially regulated coupling strength. Further supporting this hypothesis, we observe that changes in metachronal wave direction do not occur simultaneously across a band but initiate locally and propagate gradually, leading to transient coexistence of oppositely directed waves. This behavior suggests the presence of mechanical or biochemical coupling between cilia, rather than purely fluid-mediated synchronization [36, 37].

While the ciliary bands of *Didinium* support rich coordination dynamics, cells are also capable of maintaining persistent homogeneous metachronal waves leading to directed, straight swimming. In the vast majority of cells, ciliary coordination is dexioplectic, reflecting a conserved chiral bias found across a range of microswimmers [6, 11, 13]. This bias may stem from the inherent structural chirality of basal body arrangement and beat cycle asymmetries [38, 39]. Although variations in wave parameters had little impact on swimming trajectories indicating kinematic robustness, the average wavelength remained narrowly distributed and consistent across both bands and swimming modes. Notably, the wavelength closely matched cilium length, a ratio predicted to optimize propulsion efficiency in dense ciliary carpets, and was comparable to that of similarly sized spherical larvae [11, 13].

Metachronal waves are traditionally assessed primarily for their role in maximizing pumping efficiency [7, 8]. However, in the case of swimmers, the objective extends beyond simply maximizing speed, it involves efficient maneuverability for adaptive navigation and feeding strategies. Indeed, previous work has shown that experimentally observed metachronal patterns do not always align with the hydrodynamically optimal parameters predicted by theory [40, 41]. This discrepancy suggests that organisms engage in multi-objective optimization across different tasks, and that metachronal patterns are far more dynamic than commonly assumed. Indeed, our trajectory analysis of *Didinium*’s behaviors illustrate that microscale swimming is not governed solely by speed optimization, but rather by multi-objective navigation. Distinct functions such as relocation, prey detection and capture require distinct speed, lingering times and path flexibility that appear to rely on different ciliary coordination patterns and dynamics.

Together, our findings demonstrate that metachronal waves are dynamic, reconfigurable patterns capable of rapidly adjusting their symmetry, direction, and coherence to finely tune behavior. This establishes metachronal coordination not simply as a consequence of ciliary organization, but as a pivotal mechanism for behavioral flexibility. *Didinium*’s geometric simplicity and conserved ciliary architecture make it a valuable model for understanding how dynamic modulation of ciliary coordination supports complex motion typically attributed to multicellular systems. Future work elucidating the mechanisms that control ciliary coupling, trigger coordinated responses, and enable local modulation will deepen our understanding on how complex collective dynamics at the microscale encode modular behavior in unicellular protists, while informing the design of biomimetic microswimmers and synthetic cilia arrays.

## METHODS

### 1. Cell imaging

#### a. Cell culturing

*Didinium nasutum* cells were isolated from cultures purchased from Carolina Biological Supply, USA. After initial cell isolation, non axenic laboratory cultures were maintained as follows: Cells were grown at room temperature in cereal grass media (0.5 g/l cereal grass powder Fisher Scientific in Volvic water), fed with *Paramecium multimicronucleatum* twice a week and subcultured every two weeks. Before experiments, cells were removed from cultures and washed three times in Cereal grass media.

*Paramecium tetraurelia* wild-type strain d4-2 was a gift of Anne-Marie Tassin. Cells were cultured following standard protocols [42, 43].

#### b. Immunostaining and Confocal imaging

Cells were fixed and permeabilized in 4% PFA, 1% Triton X-100 in PHEM buffer for 30 minutes in room temperature. After fixation, the cells were washed with 0.05M glycine in PBS for 10 minutes and treated with blocking solution (0.2% BSA in PBS) for 15 minutes, followed by two washes in PBST (0.2% Triton X-100 in PBS) of 10 minutes each. Cells were then incubated for 1 hour at room temperature in antibody solution (0.1% BSA in PBST) with mouse anti-acetylated *α*-tubulin antibody (1:400, Sigma-Aldrich, MABT868) and rabbit anti-polyglutamate chain (Poly-E) antibody (1:1000, AdipoGen, AG-25B-0030-C050). Following three washes with PBST (10 minutes each), cells were incubated for 1 hour in room temperature with Alexa Fluor 568 goat anti-mouse IgG (Invitrogen, A11004) and Alexa Fluor 647 goat anti-rabbit IgG (Invitrogen, A27040). Cells were washed twice with PBST and once in PBS (10 minutes each) before mounted onto glass slides with VectaShield PLUS Antifade Mounting medium (Vector Laboratories, H-2000). Z-stacks of individual cells were obtained with a Leica SP8 inverted confocal laser scanning microscope equipped with a 40×/1.25 NA glycerol immersion objective.

#### c. High-speed imaging

High speed imaging of metachronal coordination and cilia dynamics were performed using either a 20x/0.8 NA air or a 40x/1.25 NA DIC water immersion objective, mounted on an Nikon Ti-E2 inverted microscope equipped with a Phantom T2540 high-speed camera (Vision research). Cells were deposited in 100 uL Cereal grass media drops on 50 mm round glass bottom dishes (Ted pella Inc, or WillCo Wells B.V.) and allowed to equilibrate for 10 minutes before imaging. Timelapses of free swimming cells of duration of 1-3 seconds were recorded at 1000-6000 frames per second.

#### d. Cell motility imaging

Live imaging of freely swimming cells was performed in custom round chambers (1.5 cm diameter, 3 mm height) laser-cut from acrylic sheets using a Glowforge laser cutter. One to three cells in cereal grass media were inserted into each chamber through a side inlet with a syringe until the chamber was full. Imaging was carried out under a dissection microscope (Leica S9E) using a Raspberry Pi HQ camera at 30 frames per second for 5 minutes per chamber.

### 2. Image analysis

#### a. Cell shape and band spacing

Z-stacks were processed in ImageJ to generate 3D maximum intensity projections and cells were reoriented so that their axis was parallel to the field of view. Cell heights were measured manually by drawing a line profile along the anterior–posterior axis excluding the cell mouth. The distance between the midpoints of the ciliary bands was also measured along the same linear profile from the Poly-E channel.

#### b. Measurement of average metachronal wave properties

A custom-made analysis pipeline was developed in Python to allow for systematic characterization of metachronal wave parameters. For each motility state, the pre- and post-processing steps were adapted to ensure accurate quantification based on the cell orientation and motion.

##### Spinning state

Timelapses were cropped to isolate individual rotating cells. Binary masks were generated by Gaussian blurring (*σ* = 20) followed by isodata thresholding with the largest connected region corresponding to the cell. Rigid registration was performed using the spam.DIC library [44], with incremental transformations computed between stepped frames (step = 0.01 s), cumulatively combined, and interpolated to yield a smooth transformation matrix. The matrix was then applied to register the timelapse and extract the simultaneous cell translation and rotation. After registration, time-lapses were normalized to correct for uneven illumination by estimating a background intensity profile using Gaussian blurring (kernel size=31) on each frame and dividing the raw image by the blurred background. Image intensities were then normalized to the range of 0 to 1.

To estimate cell size, points along the cell perimeter were manually selected on a representative frame via a graphical interface and the selected coordinates were used to fit a circle to the cell body. The obtained cell position and radius were used to define a ring-shaped mask of 22 µm height to isolate the full length of the ciliary band. The ciliary band was unwrapped using a polar coordinate transform around the detected cell center. To generate a one-dimensional representation of ciliary activity, a 3 µm thick slice through the middle of the ciliary length was averaged along the vertical axis at each position along the band, and the resulting intensities were tracked over time to produce a kymograph. To remove vertical striping artifacts, a rolling mean filter was applied along the time axis and the filtered signal was subtracted from the original kymograph.

After subtracting the mean intensity, spatiotemporal autocorrelation was computed using the Wiener–Khinchin theorem:

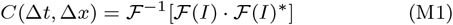

The result was normalized to *C*(0, 0) = 1. The cilia beat frequency and MW wavelength were extracted from the 2D autocorrelation by averaging the temporal and spatial power spectra, respectively, and identifying the primary spectral peaks, with associated errors estimated from their full width at half maximum (Fig. S3). MW velocity was calculated by averaging the local slopes detected in the 2D autocorrelation using the Alignment by Fourier Transform (AFT) method [45], with noise-filtered windowed analysis (window size ≈ cilia beat period, no overlap).

##### Directed swimming

Frames were Gaussian blurred (*σ* = 10) and registered using the StackReg algorithm in Python with rigid-body alignment. If necessary, images were cropped to remove black margins, and registration was repeated on the cropped stack to improve alignment. To generate the kymograph, a straight line was manually drawn perpendicular to the cell axis at the midpoint of the anterior and medial band, and the image intensity along this line was tracked over time. A rolling mean filter was applied along the time axis to suppress the vertical line artifacts originating from the cell mouth and cell body. The spatiotemporal autocorrelation was calculated using the Wiener–Khinchin theorem. Cilia beat frequency was determined from the first autocorrelation peak in time. Wavelength was typically estimated from the first spatial autocorrelation peak; however, in some cases, a double peak appeared due to perspective effects. In these cases, wavelength was corrected using formula

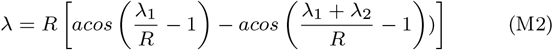

where *R* is the corresponding band radius. Metachronal wave (MW) velocity was then calculated as the product of the measured cilia beat frequency and the estimated wavelength (Fig. S3D). To correct for overestimation due to the cell spinning, surface features of the cell were tracked using CoTracker [46]. The perpendicular component of each feature track relative to the cell axis was then extracted and converted to arclengths. The spinning angular velocity (Ω) was determined as the mean slope of arclength versus time across all tracked features, with the standard deviation reported as the error. The MW parameters were then corrected using the expressions:

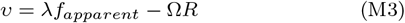

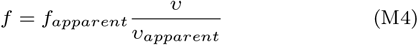

#### c. Measurement of metachronal wave dynamics

##### Phase shift asymmetries

For each rotating cell, a representative frame was isolated and the positions of fully extended cilia corresponding to the individual wave fronts were manually identified. These points were treated as individual oscillator phases to calculate the Kuramoto order pa-rameter, 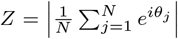 where *N* is the number of individual wavefronts along the cell perimeter and *θ*_*j*_ is the phase of the j-th individual oscillator corresponding to the position of each wavefront.. For the quantification of the degree of spatial inhomogeneity we define the order parameter as:

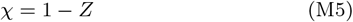

When coordination was lost for a segment of the cell perimeter, only the wavefronts at the regions where the metachronal wave remained coherent were considered.

##### Gaps in coordination

Timelapses of rotating cells where metachronal wave coordination was locally and temporarily lost were registered and kymographs were obtained following the metachronal wave analysis pipeline for spinning cells. Gaps in the ciliary wave patter appeared as noisy, non-propagating peaks interrupting the otherwise regular propagating wave pattern. To quantify the size and duration of these gaps, the local wavefront orientation was quantified using the AFT method with overlapping windows (window size = 70 pixels, overlap = 40%). Regions where the local wavefront orientation was vertically aligned corresponding to no spatial propagation (exceeded ±75°) were detected as gaps and masked (Fig. S4A). The height of the bounding box enclosing the detected region was used to measure the gap size, and event durations were defined as the time the gap width stayed above half its maximum (Fig. S4B).

##### Wave reversals and cell deflection angles

Timelapses of cells performing rapid directional reversals were registered and kymographs were obtained as in the case of the metachronal wave analysis pipeline for directed swimming. Wave reversals can be detected by changes in the sign of the local propagation slope. To minimize noise, the ciliary intensity data were first Gaussian-blurred (*σ*=5). Local wave orientations were then extracted using the AFT method (window size 50 px, 30% overlap), and the resulting angles were interpolated in first order to match the original image resolution. Wave reversal duration was quantified by tracking how much of each kymograph row maintained the original orientation. The reversal start was defined when this fraction fell below 80%, and the end when the opposite orientation occupied at least 80%. The time between these points was taken as the reversal duration. Because off-planar motion can affect the quality of this quantification, we additionally analyzed reversasl in pipette-held cells.

To estimate the corresponding deflection angles of the cell trajectory and the time it takes for the cell axis to reorient, cell masks were generated by Gaussian blurring each frame (*σ*=20) and applying isodata thresholding. The resulting binary masks were used for centroid tracking to obtain the cell trajectory and extract the instantaneous speed and cumulative reorientation (see Methods 2 d). After smoothing with a moving average filter (window size=0.01 s), cell acceleration was computed as the gradient of the speed and smoothed with a Savitzky-Golay filter (window size=0.05 s). Re-orientation duration was defined as the interval between the end of deceleration and the onset of acceleration. The end of deceleration was marked when acceleration first rose above −1 standard deviation after a period of negative acceleration, and the onset of acceleration was marked when acceleration exceeded +1 standard deviation, indicating the transition to sustained positive acceleration.The deflection angle was calculated as the difference between cumulative orientation plateaus before and after the event (Fig. S6C).

#### d. Trajectory analysis

Videos of cells swimming were pre-processed by substracting the background and applying a Gaussian filter. Individual cell tracks were obtained using the python package Trackpy. Tracks analyzed were at least 5 min in length. The tracks obtained were first smoothed using a Savitzky–Golay filter (window = 0.5 sec, polyorder = 2). The cell velocity at a given timestep was calculated as:

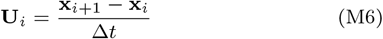

where **x**_*i*_ corresponds to the 2D coordinates of the cell position at a time *t*_*i*_ and Δ*t* is the constant time step between frames (0.033 s).

The cell orientation was calculated as:

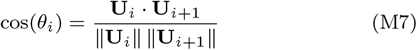

and converted to signed angles in radians ranging from *π* to +*π* using the velocity cross product. Subsequently, the angular cell velocity was calculated as:

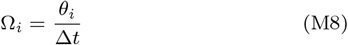

All trajectory metrics were smoothed using a centered moving average (window = 1 sec) before state classification to avoid false detections from noise. To assist in the classification the Mean Squared Displacement (MSD) and the Directional Angular Variability (DAV) were calculated over 1-second moving windows, using the expresions:

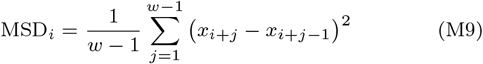

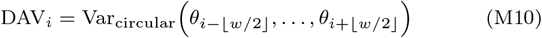

where *θ*_*j*_ is the signed angle (in radians) at point *j*, ranging from −*π* to *π*, and the circular variance was calculated using the circvar function from the SciPy library.

Spinning states were defined as segments with MSD below 5% of the mean. Directed translation segments were identified when DAV was below 50% of the mean, and all remaining segments were classified as switching states.

Velocity density maps were generated using kernel density estimates (KDEs) for each state (rotation, reorientation, forward). Filled contours were plotted for values above the 85th percentile, and labeled contours were drawn at the 85th, 90th, 95th, and 99th percentiles in the velocity (*v*) and angular velocity (Ω) space.

State-specific survival probabilities were computed using:

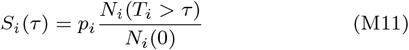

where *i* indexes the motility state, *p*_*i*_ is the fraction of total frames spent in state *i, N*_*i*_(*T*_*i*_ *> τ*) is the number of segments in state *i* with duration greater than *t*, and *N*_*i*_(0) is the total number of segments in state *i*. Survival probabilities were then fitted with the exponential decay function:

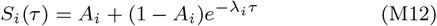

To quantify the the geometric features of helical swimming, helical segments were manually cropped from segments classified as directed motion. Helix parameters were extracted by first aligning the track using PCA, to estimate the pitch via FFT (min. frequency threshold = 0.5 *mm*^−1^ to avoid noise). To estimate radius, the aligned trajectory was divided into sliding windows (length = 1.2 times pitch, 50% overlap), and in each window, a local PCA was applied; the radius was computed by taking the average of the half of the bounding box height for each window. Helix angles were then computed as:

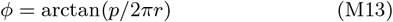

where *p* is the helix pitch and *r* the helix radius.

For the species comparative analysis, data for *P. parasitica* zoospores and *C. reinhardtii* were obtained from the published studies [31] and [32], respectively. *Paramecium tetraurelia* data was obtained in our lab by imaging and tracking cells as described for *Didinium*. Helical swimming characteristics were quantified following the same procedure as described for *Didinium*. A total of 5 cells per species was analyzed and compared. Track durations were 5 minutes for all species, except for *P. parasitica*, which were 1 minute in length.

### 3. Simulation

#### a. Equations of motion

For a spherical swimmer of radius *R* with point forces 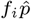 at positions 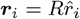, the torque is

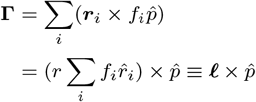

With a continuous force density *f* (*θ*), the weighted lever arm is

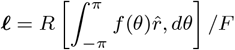

Where 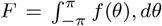 is the net propulsive force; hence |**Γ**| ≤ *RF*. Because *F* is fixed by the swimming speed, |***ℓ***| = *ℓ*_0_ is con-stant, while any azimuthal asymmetry makes ***ℓ*** precess at its own characteristic frequency. We can identify *ℓ*_0_ to 𝒳 via

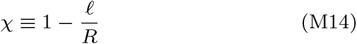

The kinematics satisfy

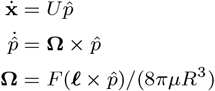

To solve this ODE exactly, we switch to a frame rotating about 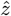 at rate *ω* so that

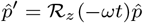

Where ℛ_*z*_ is the rotation matrix about 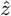. Then

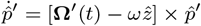

with **Ω**^*′*^ = ℛ_*z*_(−*ωt*)**Ω**. In the co-rotating frame, 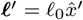 is fixed, and the angular velocity is simply 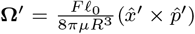. We look for solution where 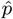 is stationary

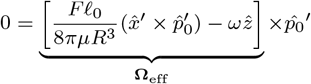

which requires 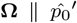. Suppose that initially 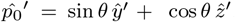, then the balancing condition is

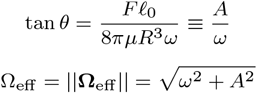

So, the helix radius is

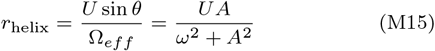

and the distance advanced per turn, or pitch, is

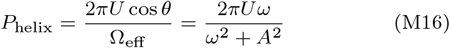

The helix andgle are related to the pitch and radius via tan *ϕ*_helix_ = *P*_helix_*/*2*πr*_helix_

#### b. Simulation details

*Didinium* is modeled as a neutrally buoyant sphere in a Stokes flow. Propulsion was produced by an array of point forces that represents individual cilia, arranged in two concentric rings of radii *R*_mid_ and *R*_ant_, separated by distance *h* = *R*. The linear density of forces are chosen to match to that of the organism, so the force per unit arc length remained constant. Force magnitudes followed a sinusoidal envelope of mean value *F* and wavelength *λ*, and oscillated at frequency *ω*. The exact magnitude is given by

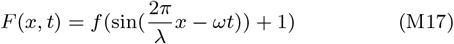

Each force vector formed an angle 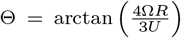 with the swimming direction 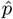, providing the tangential component required for cell rotation. Using the the experimental values of swimming speed and cell rotation speed, we estimate a net force of *F* = 1.3 × 10^−9^*N* applied at an angle Θ = 14 degrees.

During a reversal event, the point forces were flipped sequentially, implemented as a 180° rotation of an individual force about its attachment point.

Simulations and subsequent data analysis were performed in MATLAB R2024b. Cell trajectories were obtained by numerically integrating the equations of motion with the explicit Euler scheme. We repeated the simulations with progressively smaller time steps to verify that the chosen Δ*t* = 10^−3^(*R/U*) was sufficiently small and that the results were reproducible.

#### c. Randomness in wavefront distribution

Let (*x*_*i*_, *y*_*i*_) = *R* cos *θ*_*i*_, *R* sin *θ*_*i*_ for *i* = 1, …, *N*. Define the sam-ple mean 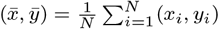 and 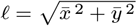.

If 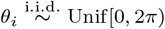 Unif[0, 2*π*), then

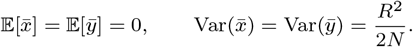

By the CLT, 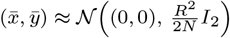, where *I*_2_ is the 2*×*2 identity matrix. So *ℓ* is approximately Rayleigh with scale 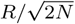 and

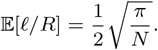

If instead the base angles *{θ*_*i*_*}* are fixed and balanced, and each point is perturbed by an independent small angular noise *ε*_*i*_ *∼* 𝒩 (0,σ^2^) *R* cos(*θ*_*i*_ + *ε*_*i*_), *R* sin(*θ*_*i*_ + *ε*_*i*_), then for σ ≪ 1,

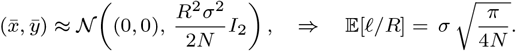

## Supplementary movie captions

**Movie 1:** Free swimming of *Didinium nasutum* shown in real time, with examples of characteristic motility states including stationary spinning, rapid reorientations, helical motion and directed straight swimming. Video was recorded using a Raspberry Pi HQ camera at 30 frames per second and preprocessed with a Gaussian filter to subtract the background.

**Movie 2:** Cilia beat cycles during directed straight swimming corresponding to the kymograph of Fig. 1B. Cilia maintain an extended configuration during the power stroke and a retracted one during off-planar recovery. Video taken with DIC microscopy and a 20x magnification objective at 1000 fps.

**Movie 3:** Free-swimming *Didinium* moving in a straight path while spinning about its axis. Metachronal waves can be seen propagating along both ciliary bands, traveling in the same direction as the cell spinning. Video taken with DIC microscopy and a 40x magnification objective at 1000 fps.

**Movie 4:** The medial ciliary band of a free-swimming *Didinium* spinning about its axis against the coverslip. Metachronal wave travels in the same direction as the cell spinning but with higher angular velocity. Cell is viewed from the posterior and cilia power strokes are towards the viewer. Metachronal waves can be seen traveling in a clockwise direction indicating dexioplectic coordination. Video taken with DIC microscopy and a 20x magnification objective at 1000 fps.

**Movie 5:** The anterior ciliary band of a free-swimming *Didinium* spinning about its axis against the coverslip (same cell as Movie 4). Cilia coordination is no longer present in this band. The metachronal waves of the medial band can be simultaneously seen as a shadow. Video taken with DIC microscopy and a 20x magnification objective at 1000 fps and registered to cancel the cell spinning.

**Movie 6:** Metachronal coordination with inhomogeneous coordination due to varying phase shift along the ciliary band. Wavelength and wave propagating velocity are larger on the left side of the circumference. Video taken with DIC microscopy and a 20x magnification objective at 1000 fps and registered to cancel the cell spinning.

**Movie 7:** Metachronal coordination with inhomogeneous coordination due to transient loss of coordination in a section of the medial band. Metachronal coordination persists for the unaffected region (top, left). Video taken with DIC microscopy and a 20x magnification objective at 1000 fps and registered to cancel the cell spinning.

**Movie 8:** Free-swimming cell switching from forward to backward swimming. The reversal of the metachronal wave pattern is visible on the anterior. The timelapse corresponds to the cell shown in Fig. 4B. Video taken with DIC microscopy and a 20x magnification objective at 1000 fps.

**Movie 9:** Movie showing a simulated metachronal wave reversal. The direction of both the cilia beat and the wave are inverted gradually along the array, following a linearly propagating front. Blue color marks the original direction and red the re-established wave.

## ACKNOWLEDGEMENTS

This work was supported by funding from the Swiss National Science Foundation Project Grant 315230 215678 to G.R.R.-S.J. A.J.T.M.M. acknowledges funding from the National Science Foundation (UPenn MRSEC DMR-2309043) and the Charles E. Kaufman Foundation (Early Investigator Research Award KA2022-129523; New Initiative Research Award KA2024-144001). The authors thank the EPFL Center for Imaging for developing an image registration code used to align videos and for their advice on image analysis.

## AUTHOR CONTRIBUTIONS

G.R.R.-S.J. designed the research. A.M.K. performed the experiments and analyzed the data. M.L. performed the simulations and developed the theoretical modeling framework under the supervision of A.J.T.M.M. The manuscript was written by A.M.K. and M.L. All authors contributed intellectually to the paper. All authors edited the manuscript.

## DATA AVAILABILITY

The data that support the plots within this paper and other findings of this study are available from the corresponding author upon reasonable request.

## CODE AVAILABILITY

The computer codes used in this paper are available from https://github.com/livingpatterns/MW_dynamics_Didinium

---

**Extended Data Table I.**
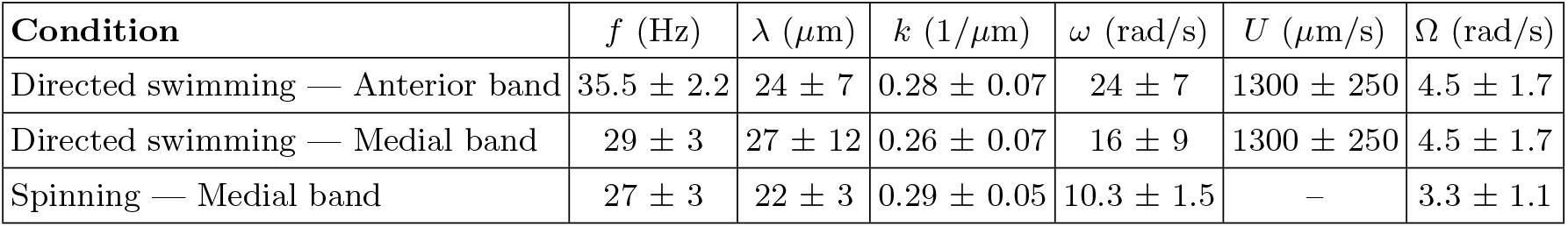
Parameters of cilia coordination and cell motility for the two bands during straight swimming and spinning in place.

**Extended Data Figure S1.**
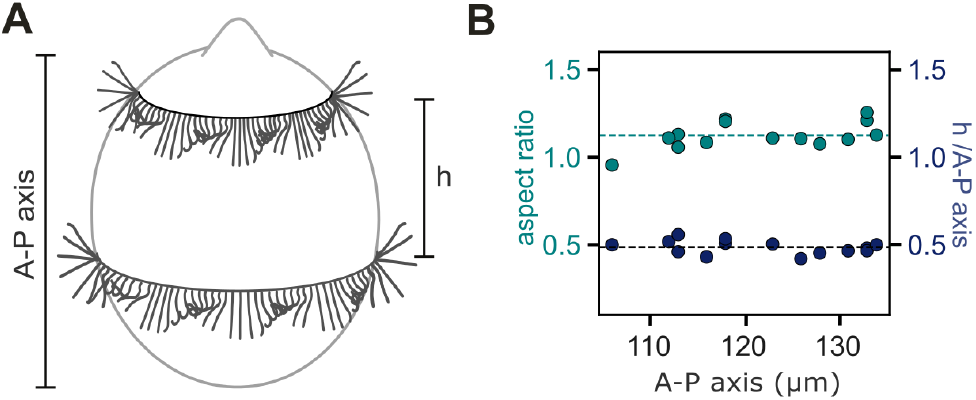
Cell proportions. (A) Schematic of a cell showing the cell height and the spacing between the two ciliary bands, *h*. (B) Aspect ratio of the cell body and band spacing, normalized by the A–P axis length, plotted as a function of cell height. *N*_*cells*_ = 14.

**Extended Data Figure S2.**
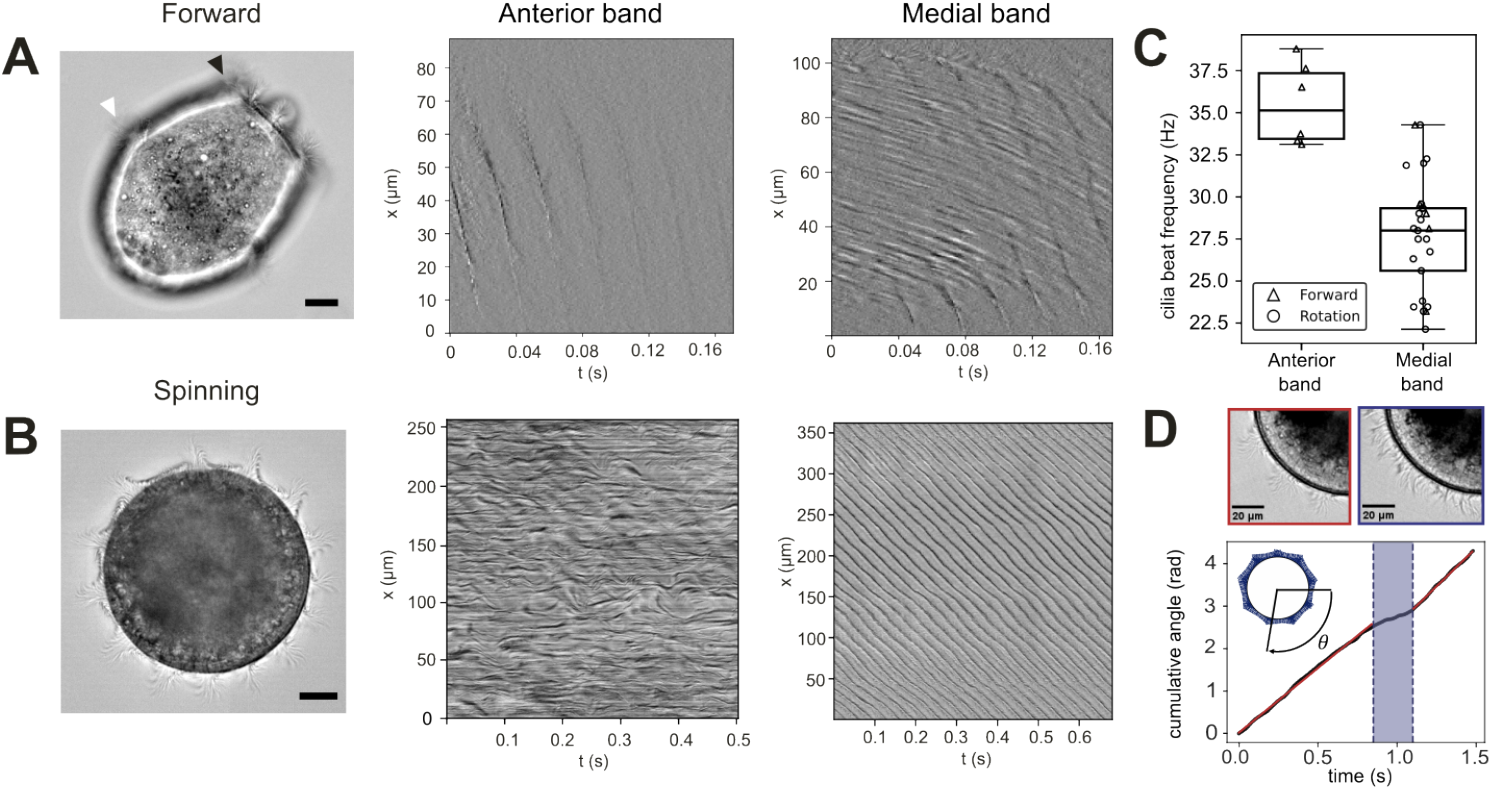
Ciliary dynamics and coordination in forward-swimming and surface-spinning cells. (A) DIC image of a free-swimming cell. Black arrowhead indicates the anterior band and white arrowhead the medial band. Right: corresponding kymographs of image intensity along the anterior and medial bands after filtering for the cell body background. Kymographs are relatively short due to limited observation time as the cells swim with high velocity, with noise from the body elements. (B) DIC image and kymographs for the two bands of a surface-spinning cell. The anterior band cilia remains uncoordinated during this motion. (C) Boxplots of ciliary beat frequency measurements for both bands and motility states. Sample sizes: *N*_*f*_ = 6 forward-swimming cells, *N*_*s*_ = 16 surface-spinning cells. (D) Top: Segment of the cell showing regions of preserved coordination (red window) and temporary local loss of coordination (blue window). Bottom: cumulative turning angle over time. Red lines indicate linear fits before and after the loss of coordination, with shaded blue area marking the event. Fit parameters: Ω_before_ = 3.048 *±* 0.015 rad*/*s, Ω_after_ = 3.627 *±* 0.024 rad*/*s.

**Extended Data Figure S3.**
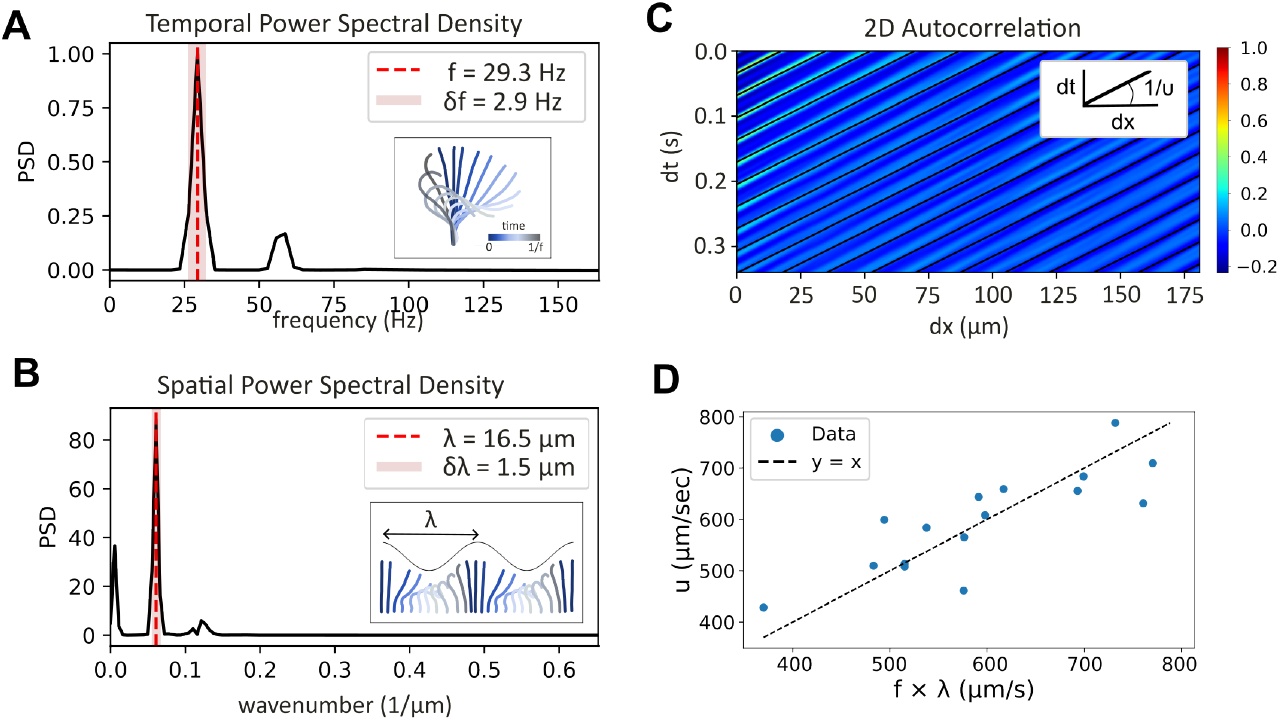
Quantification of MW parameters. (A) Temporal power spectral density (PSD) of the signal. The dominant frequency peak (red dashed line) is used to extract the ciliary beat frequency, with the shaded region denoting its error. Inset: schematic of ciliary beat cycle. (B) Spatial PSD of the MW pattern. The dominant wavelength peak (red dashed line) is used to extract the wavelength, with the shaded region representing the error. Inset: schematic of a MW marking the wavelength. (C) Two-dimensional autocorrelation of the MW pattern. Black diagonal lines mark the detected slope, which corresponds to the wave velocity. (D) Comparison between the measured wave velocity, *u*, and the product of measured frequency and wavelength, *f × λ*, showing good alignment with the *y* = *x* (dashed line). In the case of forward swimming velocities were thus calculated with this expression.

**Extended Data Figure S4.**
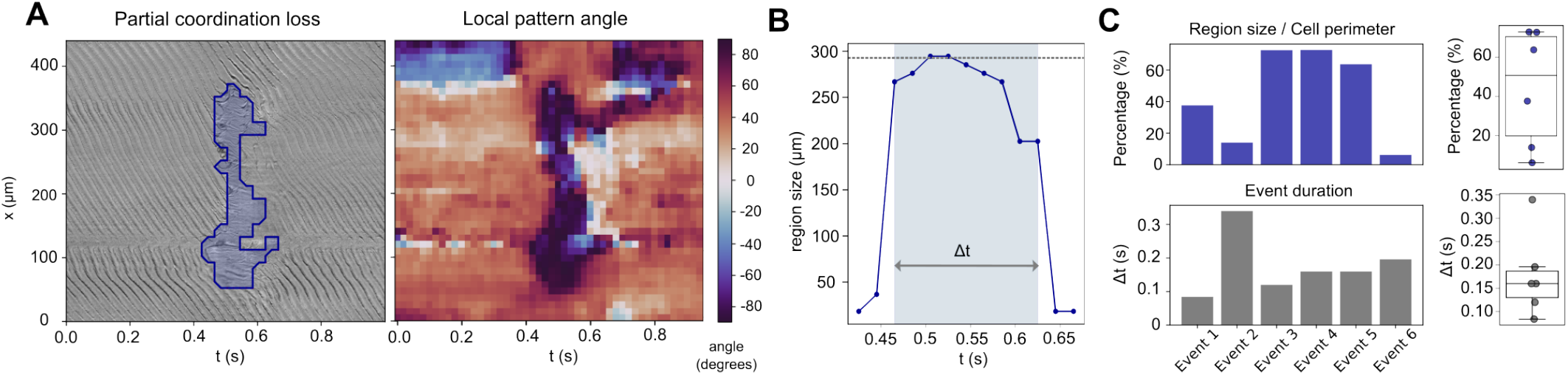
Quantification of transient loss of coordination. (A) Kymograph of image intensity in a medial band during a representative event of partial loss of coordination (left). The blue shading marks the automatically detected area of the affected region. Right: corresponding colormap of local orientation angles in the same region. The region shown in A corresponds to detected angles with absolute values ≥ 75°. (B) Time evolution of the affected region size of the event of panel A. The dotted line marks the maximum region size during the event. The event duration, Δ*t*, is defined as the time span for which the region size exceeds half of its maximum value. (C) Quantification of individual events. Left: bar plots of affected region size normalized to cell perimeter (top) and event duration (bottom). Right: corresponding boxplots.

**Extended Data Figure S5.**
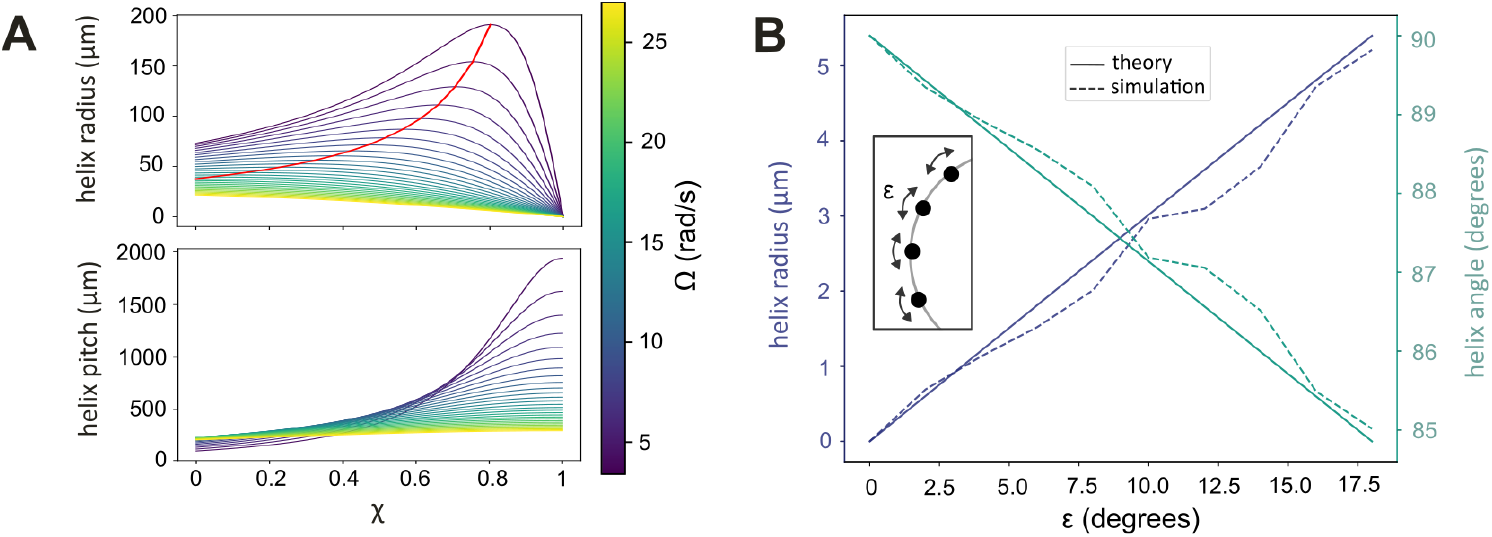
Helix geometrical characteristics under heterogeneous force distribution. (A) Dependence of helix radius (top) and helix pitch (bottom) on the heterogeneity parameter, 𝒳, for different values of Ω. Solid lines represent theoretical predictions. Red line indicates the global maxima of the helix radius. (B) Influence of Gaussian noise in an otherwise homogeneous force distribution on helix radius and helix angle. Solid lines indicate theoretical predictions, while dashed lines show simulation results. The inset illustrates representative wavefront placements (black dots) with arrows denoting noise amplitude.

**Extended Data Figure S6.**
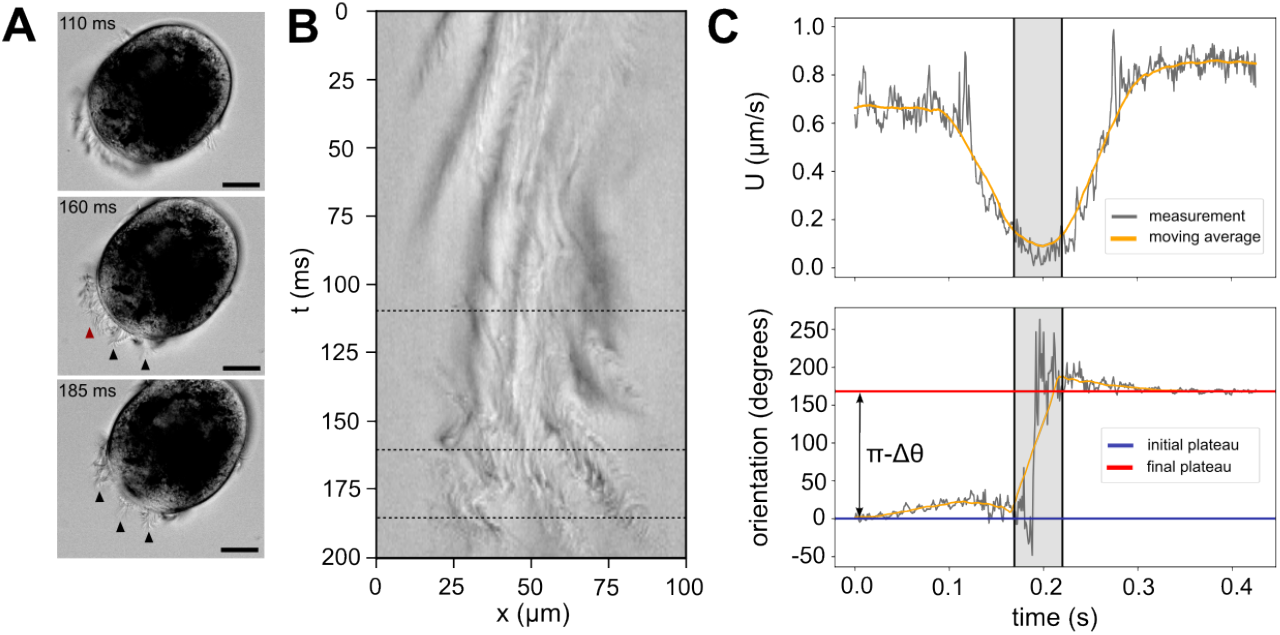
MW dynamics and cell deflection during cell reorientation. (A) Snapshots of a cell undergoing direction reversal showing the progression of the MW pattern re-emergence. Black arrows mark the new wavecrests, while the red arrow highlights the region that remains uncoordinated. (B) Kymograph of image intensity along the anterior band, with dashed lines indicating the timesteps corresponding to the snapshots in A. (C) Motility analysis of direction reversals. Data correspond to a different cell than the one in A-B. Top: cell velocity as a function of time. Bottom: cumulative cell orientation over time. Grey lines show raw measurements, yellow lines the smoothed signal obtained via moving average, and shaded areas mark the reversal duration. The initial (blue) and final (red) orientation are marked with horizontal lines.

